# Diagnostic and Therapeutic Microbial Circuit with Application to Intestinal Inflammation

**DOI:** 10.1101/2020.11.10.377085

**Authors:** Liana N. Merk, Andrey S. Shur, Smrutiti Jena, Javier Munoz, Douglas K. Brubaker, Richard M. Murray, Leopold N. Green

## Abstract

Bacteria genetically engineered to execute defined therapeutic and diagnostic functions in physiological settings can be applied to colonize the human microbiome, providing *in situ* surveillance and conditional disease modulation. However, many engineered microbes can only respond to single-input environmental factors, limiting their tunability, precision, and effectiveness as living diagnostic and therapeutic systems. For engineering microbes to improve complex chronic disorders such as inflammatory bowel disease, the bacteria must respond to combinations of stimuli in the proper context and time. This work implements a previously characterized split activator AND logic gate in the probiotic *Escherichia coli* strain Nissle 1917. Our system can respond to two input signals: the inflammatory biomarker tetrathionate and a second input signal, anhydrotetracycline (aTc), for manual control. We report 4-6 fold induction with a minimal leak when the two chemical signals are present. We model the AND gate dynamics using chemical reaction networks and tune parameters *in silico* to identify critical perturbations that affect our circuit’s selectivity. Finally, we engineer the optimized AND gate to secrete a therapeutic anti-inflammatory cytokine IL-22 using the hemolysin secretion pathway in the probiotic *E. coli* strain. We used a germ-free transwell model of the human gut epithelium to show that our engineering bacteria produce similar host cytokine responses compared to pure cytokine. Our study presents a scalable workflow to engineer cytokine-secreting microbes. It demonstrates the feasibility of IL-22 derived from probiotic *E. coli* Nissle with minimal off-target effects in a gut epithelial context.

## 1 Introduction

The global burden of chronic disease is quickly becoming a universal crisis, with one in three adults suffering from multiple chronic conditions [1]. While the exact causes of chronic diseases remain elusive, comprehensive studies attribute abnormal interaction between the epithelial immune response and the resident microbiome as a primary cause [2]. Approximately 10^13^–10^14^ bacterial cells live in the dynamic and complex community within the gut microbiome, where they can impact numerous facets of human health [3–5]. The hypothesis that changes in the composition of the microbial community result in disease offers a compelling motivation for engineering microbes to sense, treat, and prevent chronic conditions. Engineering microbes for diagnostics and therapeutics is a growing field in synthetic biology, owing to the tractability and relative safety of genome engineering in microbes [6–8]. Due to the presence of bacteria in various ecological niches, many evolved sensors and effectors exist for therapeutically relevant molecules [9–12]. Microbes’ ability to sense and respond to stimuli *in situ* offers controlled and targeted responses to pathogen challenges or chronic inflammation in areas of the body that are traditionally difficult to access.

Microbes capable of drug manufacture and secretion could colonize the microbiome while providing long-lasting *in situ* therapeutics without complex dosing schedules. However, these microbes express target molecules constitutively, not taking advantage of the ability of bacteria to sense and respond to their environment. Furthermore, engineered signal processors in bacteria are motivated by a need to interpret complex immune signaling environments, particularly in conditions that result in chronic disease. To address this need, logic gates have been constructed and characterized in bacterial and mammalian cells [13,14].

Synthetic gene circuits have demonstrated digital logic function but often lack modularity. *Escherichia coli’s* modular cloning library of promoters and coding sequences has allowed for the programmable sensing of an inflammatory signal tetrathionate [15]. Single-element decision-making has limited robustness due to biological noise and lack of environmental specificity. However, integrating two or more signal inputs can improve detection precision by pattern recognition. Recent work has demonstrated the advantage of detecting multiple inputs versus a single input in *E. coli*, where synthetic genetic counting networks can precisely determine phasic signal patterns [16]. Notably, the split activator AND gate in *E. coli* has been constructed using an orthogonal regulatory component from the hypersensitive response and pathogenicity (hrp) pathway derived from *Pseudomonas syringae* [17]. The modularity of the hrp logic system presents opportunities to couple various exogenous inputs (i.e., pro-inflammatory markers) to develop more complex molecular logic gates such as AND, NOR, NAND, and XOR systems [18].

Logic-gated outputs allow a circuit to respond to complex microbiome perturbations such as inflammation, which can drive dysbiosis in microbial populations [19]. Dysbiosis is a root cause of inflammatory bowel disease (IBD) and has been associated with infections, obesity, and other medical challenges [20]. Currently, IBD’s most common treatment is large doses of oral anti-inflammatory drugs, which have broad and nonspecific effects that do not account for local environmental changes [21]. These anti-inflammatory drugs often require irregular and frequent dosage schedules that are challenging to adhere to, with the average patient missing half of their treatments [22].

Recent studies have found promise for inflammation treatment with microbes secreting immunomodulators such as AvCystatin [23], anti-tumor necrosis factor alpha [24], interleukin-27 (IL-27) [25], and IL-10 to maintain barrier tissue integrity during pathogen invasion [26]. Interleukin 22 is a member of the IL-10 cytokine family and is expressed by innate and adaptive immune cells [27]. Although the function of IL-22 in intestinal homeostasis remains controversial [28], it is thought to coordinate innate defense mechanisms such as mucus production and epithelial regeneration [29]. Due to its integral role, IL-22 is a good candidate as a therapeutic against IBD.

While microbes have been engineered to respond to environmental factors like inflammation, they are often limited to sensing single inputs. In this work, we develop a logical response circuit in *E. coli* Nissle that secretes mammalian IL-22 in the presence of the inflammation-associated biomarker tetrathionate and an external activator isopropylthiogalactoside (IPTG) or anhydrotetracycline (aTc) [30]. We build upon the previously characterized hypersensitive response and pathogenicity (hrp) split activator AND logic gate reported by Wang et al. [17]. We develop a mathematical model of the tetrathionate two-component system to explore the design space of our circuit components and characterize the effects of leak. Here, we introduce an engineered microbial prototype that processes multiple inputs to respond to inflammatory environments.

## 2 Results and Discussion

### 2.1 Tetrathionate sensor validation

Inflammation is a critical component in chronic conditions, including IBD. Engineering bacterial circuits capable of sensing inflammation-associated biomarkers have been highlighted in recent synthetic biology approaches [31]. Gut inflammation induced by mucosal microbes generates reactive oxygen species, resulting in the metabolite tetrathionate [32]. Daeffler et al. identified a tetrathionate two-component sensor from the marine bacterium *Shewanella baltica* OS195 [9]. The system comprises a membrane-bound sensor histidine kinase, TtrS, and a cytoplasmic response regulator, TtrR. Tetrathionate binds to TtrS, causing phosphorylation, leading to a complex that can phosphorylate TtrR. Phosphorylated TtrR, in turn, activates the promoter P_Ttr_, which demonstrates low cross-activation by a range of other ligands likely present in the gut.

We incorporate this previously characterized two-component tetrathionate detection system into an engineered *E. coli* chassis as a therapeutic strategy for microbiome applications. A critical design constraint is to minimize the number of plasmids in the organism. Using 3G assembly [33], we constructed a single construct expressing the necessary regulators and the tetrathionate inducible promoter, P_Ttr_, shown in Figure 2A. To minimize leak caused by polymerase readthrough, we incorporated the inducible promoter upstream from the constitutively expressed regulators. We optimized the regulator expression level by varying the ribosome binding sequence RBS preceding both tetrathionate sensing regulators TtrS and TtrR using the Andersen RBS pool [34]. Six successful constructs were sequenced, and RBS strengths were estimated, as shown in Figure 2B [35]. Weak RBS preceding TtrS or TtrR leads to a lower fold change of GFP expression upon the addition of tetrathionate, as seen in the simulated results in Figure 2C. The high-activation circuit construct was selected for future experimental builds due to 13.8x activation with maximum tetrathionate input.

**Figure 1.**
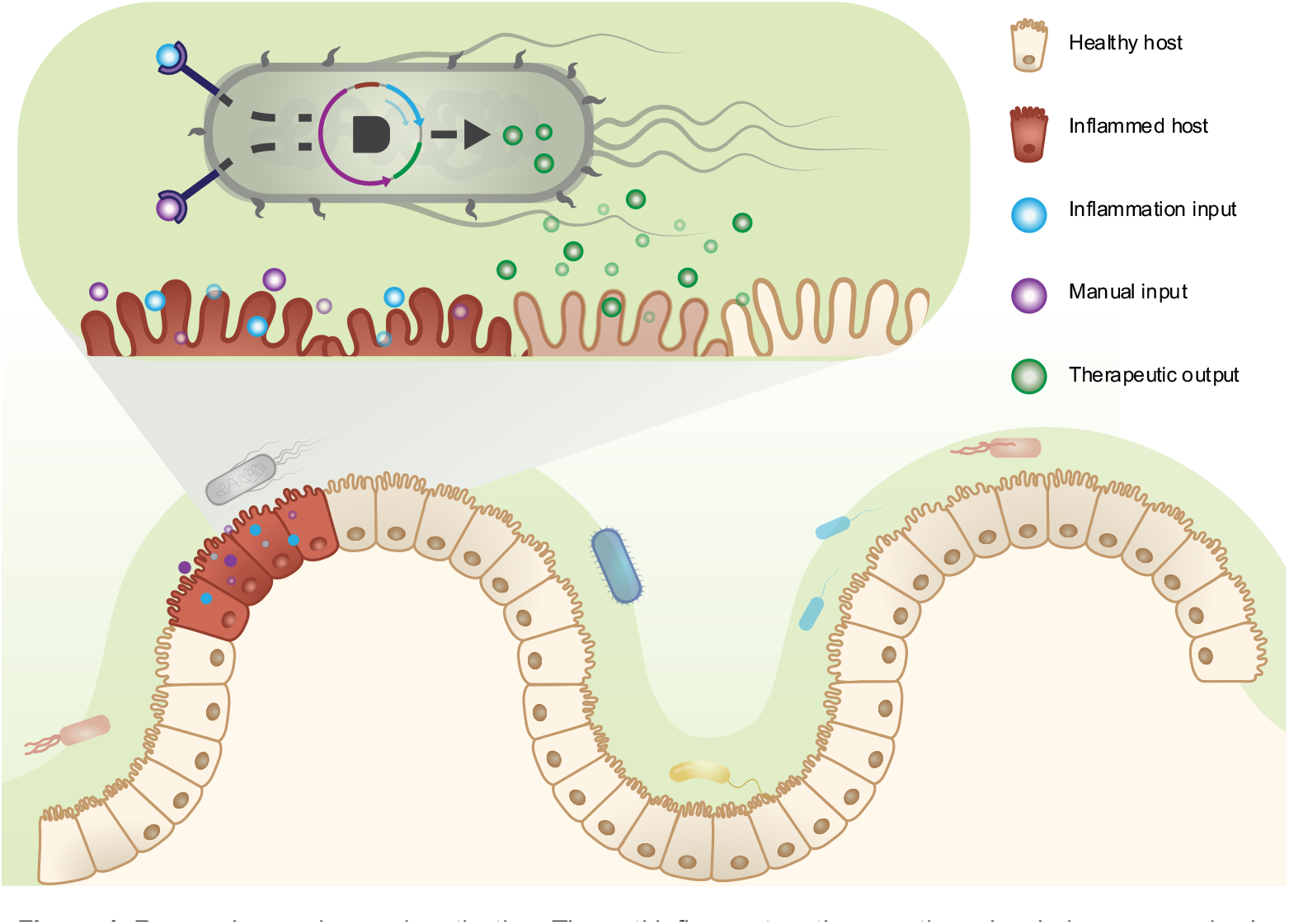
Research overview and motivation. The anti-inflammatory therapeutic molecule is expressed only when the genetically engineered bacterial cells detect the inflammatory biomarker and manual control inputs, minimizing off-target effects. Figure not drawn to scale.

**Figure 2.**
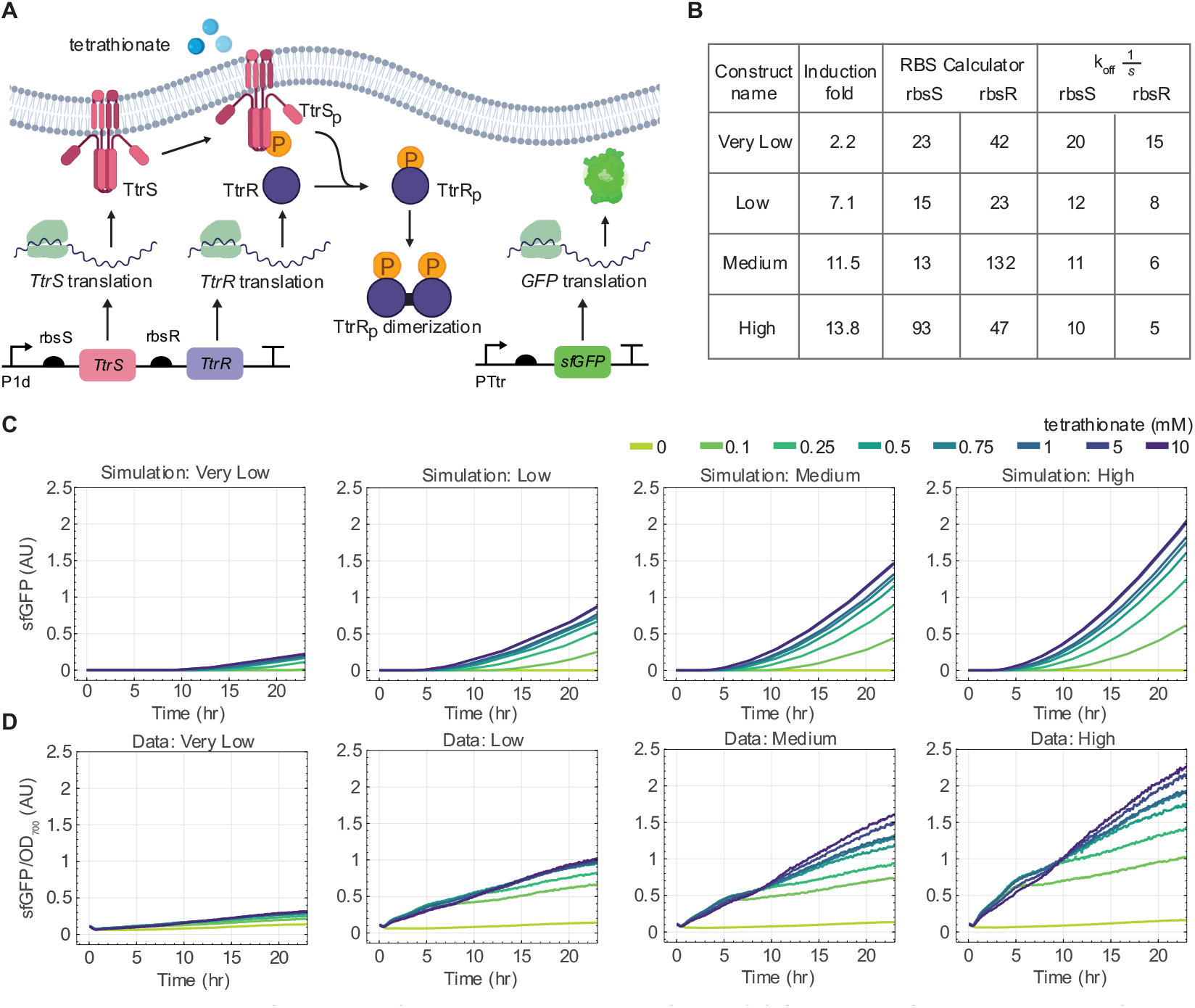
Expression and Simulation of Tetrathionate Response Circuit, (A) Schematic of Tetrathionate Two-Component System. All shown steps were modeled using chemical reaction networks. (B) Relative RBS binding strengths toTtrS and TtrR were calculated from sequenced RBS sites isolated from 4 functioning tetrathionate response circuits. Off rates for simulated reactions. (C) Simulations of RBS tuning by varying ribosome off rates to each ribosome binding site. (D) Plate reader time course with increasing tetrathionate concentrations and different ribosome bindingstrengths.

### 2.2 Modeling the Tetrathionate Two-component System

Two component signaling is one of the most prevalent transmembrane signal transduction and gene expression pathways in microbes [36]. Therefore, we derived a mathematical model from chemical reaction networks to describe the dynamics of the molecular signal transduction pathways. This aided our understanding of various experimental optimization measures, such as the ribosome binding strength tuning of the fluorescent output. Our model is divided into three parts: (1) the expression of tetrathionate regulators TtrS and TtrR, (2) the phosphorylation of the tetrathionate regulators, and (3) the reporter gene’s production.

The regulator’s TtrS and TtrR constitutive expression is under the common promoter J23103 (P1d) [34]. Since these parts are co-transcribed, we model a single transcription reaction for *TtrS* and *TtrR* gene regulators. Transcription is modeled as a two-step process: (1) binding the promoter to the RNA polymerase to form a complex and (2) transcription of the promoter-RNA polymerase complex to produce the corresponding mRNAs. Since each regulator controls its ribosome binding site, we modeled two independent translation processes of *TtrS* and *TtrR*. Furthermore, we modeled translation as a two-step process where (1) the ribosome binds to the mRNA transcript to form a complex that then (2) translates to the production of the regulator protein components.

For the phosphorylation pathway, we model the tetrathionate molecule binding to TtrS as a reversible reaction when bound, initiating the phosphorylation of TtrS. The cytoplasmic response protein TtrR can bind to the phosphorylated or dephosphorylated TtrS. If TtrR binds to dephosphorylated TtrS, there is a higher OFF rate than when TtrR binds to phosphorylated TtrS. Phosphorylation of TtrR only occurs when TtrR binds to the phosphorylated TtrS, forming a complex, TtrR: TtrS^P^. It is unknown whether the dephosphorylation of TtrR is phosphatase-dependent, like the KdpD/KdpE TCS in *E. coli* [37]. Here, we do not model phosphatase explicitly but rather set an explicit rate, *k*_*dephos*_, that defines the dephosphorylation of TtrR.

The third and final part of the two-component signaling model is the activation and expression of the GFP reporter gene. Phosphorylated TtrR dimerizes within the cytoplasm, which we model as a reversible reaction. It is unknown whether inactive TtrR (dephosphorylated dimers or monomers) can bind to the promoter region. Here, we only model the dimerized, phosphorylated TtrR as activators of downstream gene expression. Once the activated TtrR binds, the inactive P_Ttr_ promoter is converted to an active state, denoted as PTtr^*^. In our model, RNA polymerase can only bind to the activated promoter PTtr^*^. The RNA polymerase:P^Ttr*^ complex triggers the transcription of the GFP mRNA, GFP_T_. The GFP transcript reversibly binds with a ribosome and is irreversibly converted into the unbound ribosome, GFP_T_, and GFP protein.

Choosing the proper reaction rate parameter values for each chemical reaction is critical in developing a signal transduction model. Since finding exact values for all the rate parameters *a priori* is difficult, we make a few simplifying assumptions. We assume the explicit translation rates are identical for all mRNAs in the system. Furthermore, we assume that all degradation reactions occur at the same rate. We selected the nominal values from the various results and information available in the literature, discussed further in the supplementary information for all parameters. Since the ribosome binding site (RBS) strength is a tunable parameter in the experimental design, we keep this parameter open to change during the simulations to observe the potential effects of RBS sequence strengths. The different RBS strengths are modeled by changing the ribosome’s unbinding reaction rates to a given transcript.

The model simulations with varying ribosome binding strengths are shown in Figure 2C. The model predicts that the RBS preceding the inducers strongly affects output fold change. The experimental results follow similar trends to the simulated RBS tuning results, as shown in Figure 2D. The detailed model, including the system’s species, reactions, and parameters, is given in the supplemental information, with all code publicly available on GitHub [link].

A future line of work would be to quantitatively validate the model parameters by fitting the experimental data to the simulations so that the model can be used to make credible predictions on future circuit designs and expression dynamics. Since various parameters in the model are context-dependent, parameter tuning of a validated model *in silico* may provide helpful insights when implementing this circuit in complex, *in vivo*, such as designing responsive therapeutic circuits for the gut. Similarly, the effects of resource sharing and high burden due to the expression of proteins may be quantified using this model.

### 2.3 Engineering two-input AND Gate

To incorporate Boolean AND logical sensing, we implemented a co-dependent split activator system that consists of the HrpR and HrpS regulators to drive the expression of our GFP reporter gene. We coupled our previously optimized inflammation tetrathionate-dependent promoter to drive the expression of the HrpR regulator. The second regulator, HrpS, is driven by P_Lac_ activated by the chemical inducer IPTG. The HrpR and HrpS split regulators form a homo-hexameric complex needed to activate P_HrpL_, which drives GFP expression, shown in Figure 3A. Thus, the engineered logical AND gate will express the GFP output only in the presence of tetrathionate and IPTG, following the logic table in Figure S3B.

**Figure 3.**
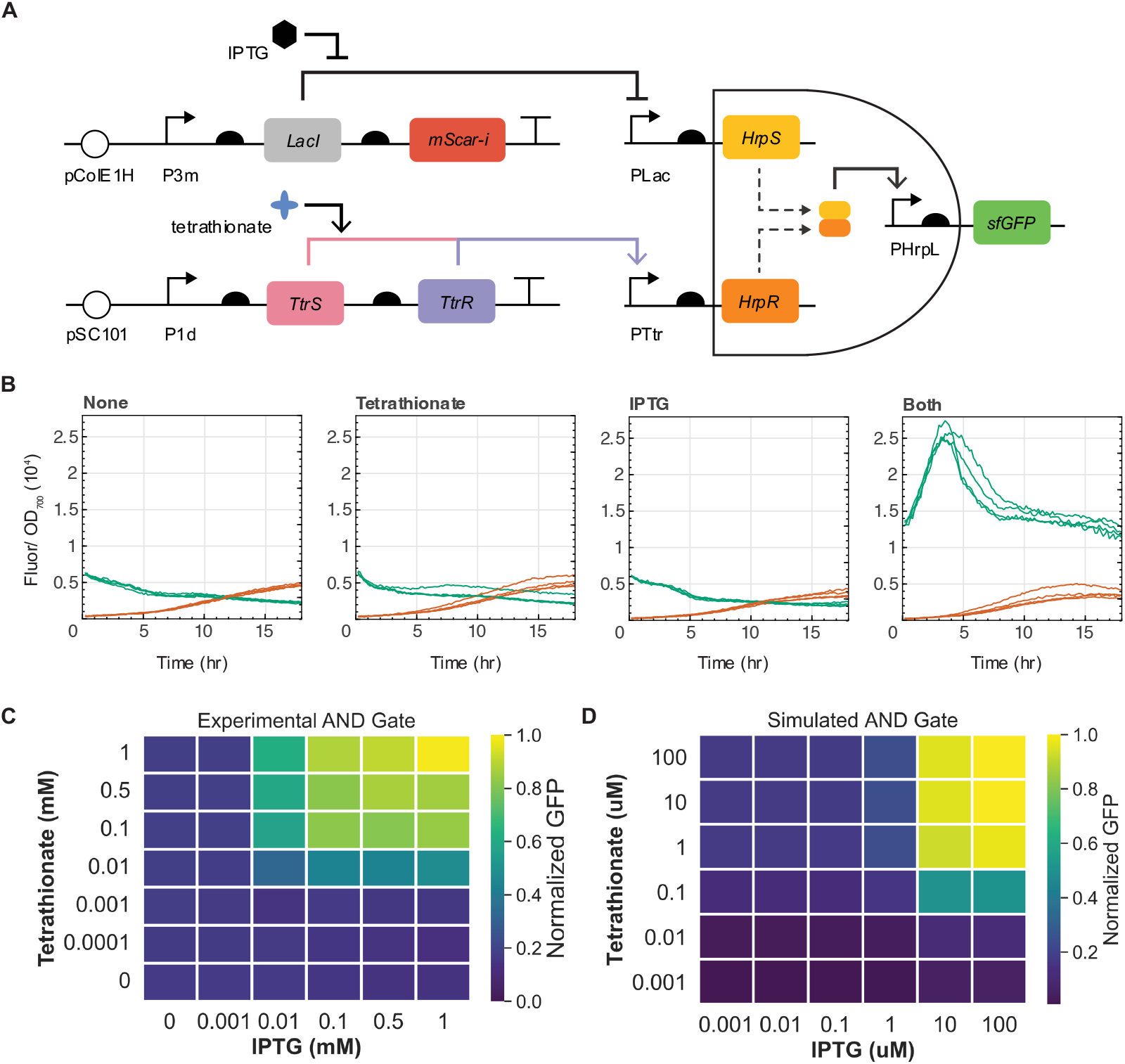
AND Gate Design and Screening. (A) Schematic of engineered logic-based theranostic: We expect GFP expression only when both inducers are present. (B) Plate reader assay for Nissie AND constructs. Maximum induction of 1 mM tetrathionate and 1 mM IPTG was used. Fluorescence values were normalized to OD700 readings and readings shown begin at log-growth phase. (C) Plate reader assay for grid of inducer concentrations. Values displayed are GFP fluorescence normalized to OD, divided by the maximal value, achieved at 1 mM IPTG and 1 mM tetrathionate. (D) Heatmap of simulated AND gate. We model the expression of the two split activator components, describing all chemical reactions in Table S1 with parameters in Table S2. Note the inducers are different orders of magnitude because our model functions in bulk cell extract rather than cells and thus are only relatively, not absolutely, comparable to concentrations in 3C.

While this split activator system has previously been tested in seven different chassis [17], it has not been tested in *E. coli* Nissle 1917. We first performed one round of optimization of the AND gate components in *E. coli* Marionette Clo cells [40], which includes the genome-integrated expression of LacI. We initially used the RBS sequences B0034 for HrpR and rbsH for HrpS reported by Wang et al. [17]. However, the expression was leaky in all inducer conditions, suggesting that the activator expression was too high. We cloned the Anderson library of ribosome binding sites (ARL) [34] upstream of the HrpR regulator to resolve the undesired leak. We then isolated five constructs with significant fold change in response to both inducers (IPTG and tetrathionate). Constructs were isolated, sequence verified, and re-transformed into Nissle, which required optimization of LacI expression levels. Using a stereoscope for visual screening, we identified colonies with functioning dual-plasmid AND logic gate construct (Figure S3D). Our experimental results from the plate reader fluorescence illustrate a 6-fold induction in both tetrathionate and IPTG inducers and minimal expression in no inducer or single-induction conditions (Figure 3B). We further show that the AND gate displays digital-like activation across a range of both inputs Figure 3C. To our knowledge, this is the first engineered inflammation-responsive AND logic gate in *E. coli* Nissle.

### 2.4 Secreting Anti-Inflammatory Molecules

Bacteria have various secretion pathways for transporting proteins from the cytoplasm into microcompartments and to the environment, coordinating cell-to-cell interactions [41–43]. One such system is gram-negative bacteria’s type I secretion system (T1SS). Specifically, the hemolysin secretion system is released from pathogenic bacteria through a transmembrane pore previously engineered to secrete other metabolites in *E. coli*.

Here, we construct a plasmid containing the HlyB (inner membrane) and HlyD (transmembrane) sequences. TolC, the outer membrane protein, is present in the *E. coli* genome. We hypothesized that the stoichiometry of HlyB and HlyD is vital for secretion efficiency; thus, we used the ARL RBS pool preceding the *HlyB* and *HlyD* genes to screen for optimal expression levels of each secretion component. First, constructs containing P_Lac_-inducible human IL-10 or mouse IL-22 were assembled onto the pColE1 high copy number backbone. These plasmids were sequence-verified and co-transformed into *E. coli* HB2151 with the constitutive *HlyBD* plasmid, as shown in Figure 4A. Optimization for the maximal secretion of IL-22 is described in the Supplemental figures S4-S6. Through multiple rounds of optimization, IL-10 secretion constructs were unsuccessful (i.e. no signal in the supernatant), perhaps indicating an incompatibility with IL-10 and this particular secretion mechanism. While it has been shown that the size of the product does not affect the secretion [46], the folding rate can impede the secretion if folding occurs intracellularly. Thus, it may be the case that mouse IL-10 folds faster than the hemosylin machinery can export the cytokine. An alternative strategy for secreting IL-10 from the engineered bacterial chassis is incorporating inducible lysis constructs such as the well-characterized PhiX174 bacteriophage protein E lysis protein [47,48].

**Figure 4.**
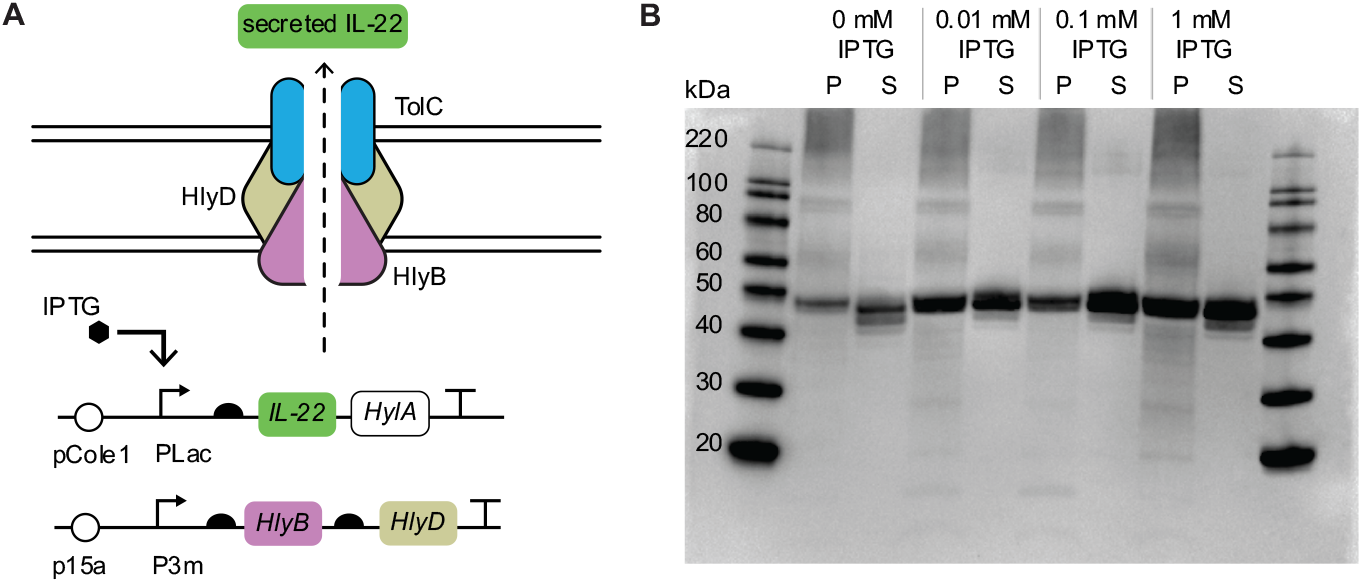
High Yield IL-22 E. coli Nissle. (A) Constructs with constitutively expressed secretion machinery hlyB and hlyD located on the p15 backbone and IPTG-inducible hlyA and His-tagged IL-22 located on the ColE1 high copy backbone. (B) A western blot with anti-His antibodies showed a strong band signifying IL-22 secretion. Band strength increases with increasing induction of IL-22 expression. (P: pellet. S: supernatant.)

For IL-22, the expected band size for the HlyA-tagged IL-22 monomer is 46 kDa. We see this band present in all inducer conditions in 4B, indicating insufficient repression of P_Lac_ in the cell. This is expected, as we chose a high expression RBS preceding IL-22 and a high copy number plasmid for this initial secretion optimization in the event the section band would be faint. However, because the final construct would be under the control of the optimized AND gate, we did not continue to optimize this construct. We mini-prepped this successful HlyB-D secretion plasmid, antibiotic selected to isolate only secretion, then created the AND-inducible secretion construct.

### 2.5 AND Secretion of IL-22

The excessive nature of inflammatory mediators in chronic intestinal immune response calls for designing living therapeutics that integrate multi-input diagnostics with control systems to reduce prolonged inflammation-associated conditions. An ideal diagnostic-therapeutic system can multiplex environment signals to decide the dosage and timing of the therapeutic output. Here, we couple the AND logic gate and tune genetic components to drive the logical secretion of mouse IL-22 using probiotic *E. coli* Nissle (Figure 5).

**Figure 5.**
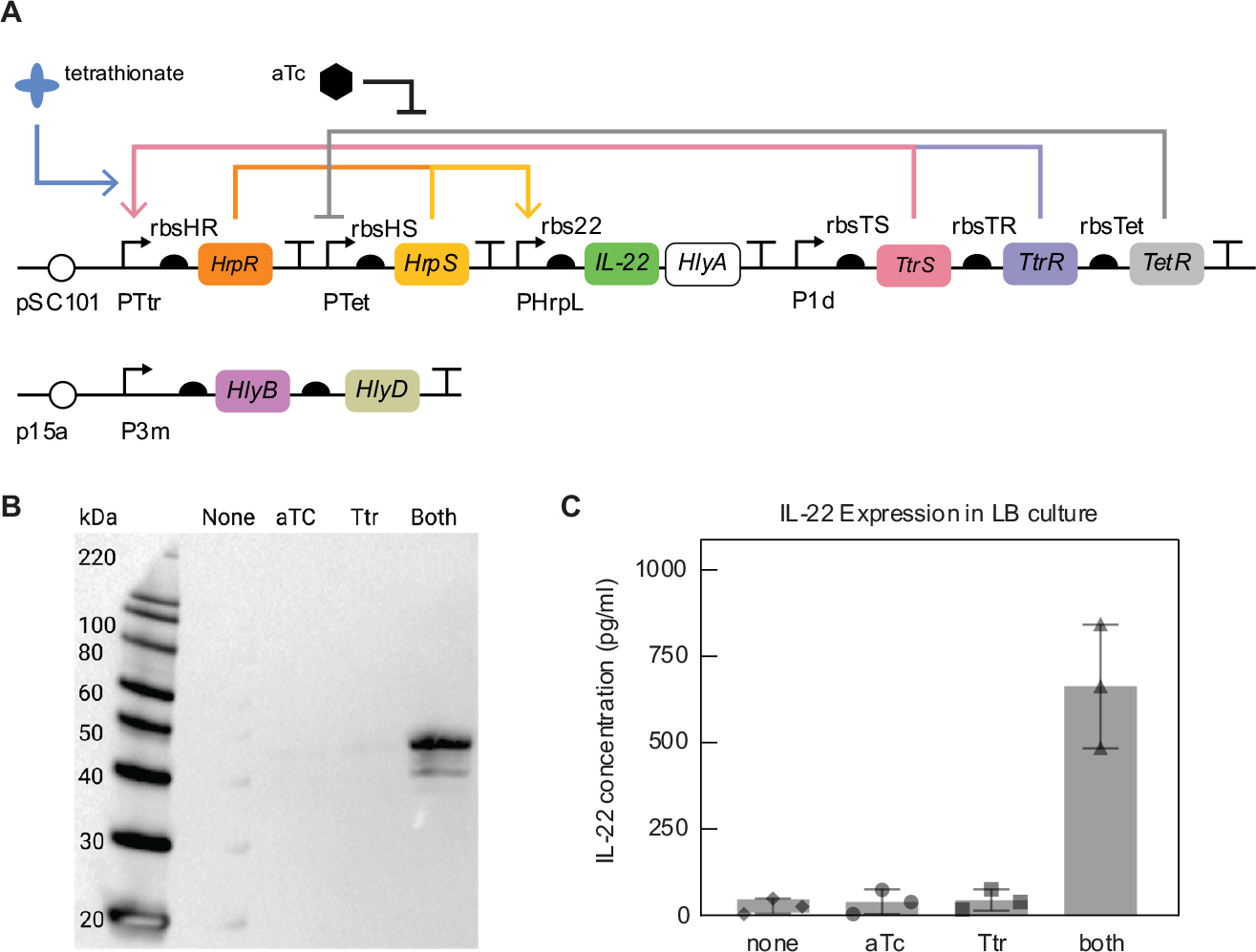
AND Gate IL-22 Secretion. (APlasmid diagram of the logic gate inflammation diagnostic components to drive the production of the HlyA-tagged IL-22 secretion system. (B) A western blot with anti-His antibodies shows IL-22 secretion only when aTc and tetrathionate are present. (C) ELISA quantifying the secretion of IL-22 in response to the logic induction of both tetrathionate and aTc inducers after 6, 12, and 24 hours of inoculation.

Recall that the initial AND logic gate shown in Figure 3B was optimized with *LacI* and *mScarlet-i* components on a separate plasmid from the logic gate components. As we include the secretion machinery components, we aim to reduce the final plasmid load on our cells. Thus, we consolidated the AND gate, repressors, and mScarlet-i onto a single plasmid and re-optimized. During this optimization, we were able to get less leaky performance using the aTc-inducible P_Tet_ promoter in place of the IPTG-inducible P_Lac_ promoter. Once we had our functional single-construct AND gate, we replaced the AND-activated GFP with the IL-22-HlyA construct. We decided to keep a relatively strong UTR1 as the RBS preceding IL-22 to allay concerns that the signal would be too weak to detect. We co-transformed the previously validated constitutive secretion machinery plasmid with AND-IL-22-HlyA plasmid.

We tested two sequence-verified colonies, finding one showing the correct size band for secreted IL-22 only in the presence of both inducers, shown in Figure 5B. Interestingly, there does not appear to be a band in the pellet of both inducer conditions (S6B), potentially signifying that the production rate matches the secretion rate now that the construct is on a lower copy-number plasmid. Finally, we quantify the concentration of IL-22 secreted using ELISA assay in varying inducer conditions after 6 hours, 12 hours, and 24 hours inoculation in LB at 37°C. Figure 5C illustrates that the logical secretion of IL-22 reaches a concentration of 700 pg/ml after 6-hour growth. Culture inoculation for 12 and 24 hours results in leaky expression of IL-22 in conditions without inducers or signal inducer conditions. This is likely due to non-ideal growth dynamics as the engineered bacteria are no longer in the exponential-growth phase after 12 hours of inoculation but in the lag phase, where high cell lysis rates would release non-secreted IL-22 into the supernatant.

### 2.6 Biological Activity of Microbe-Produced IL-22 in a Human Germ-Free Gut Model

Characterizing the biological effects of the bacterial-produced cytokine compared to the pure cytokine is essential in assessing the potential translational value of an IL-22-secreting microbial therapeutic. To this end, we co-cultured Caco2 and HT-29 colorectal cancer cells on transwell inserts to create a germ-free intestinal epithelium model. This germ-free transwell model mimics critical aspects of the human intestine, such as tight junctions’ expression, mucins’ production, and cytokine mediators [49,50]. Using this platform, we assessed the comparative effects of pure IL-22 and microbe-produced IL-22 and associated microbial products on modulating intestinal epithelial cytokine responses in the presence of a lipopolysaccharide (LPS) pro-inflammatory stimulus (Figure 6).

**Figure 6.**
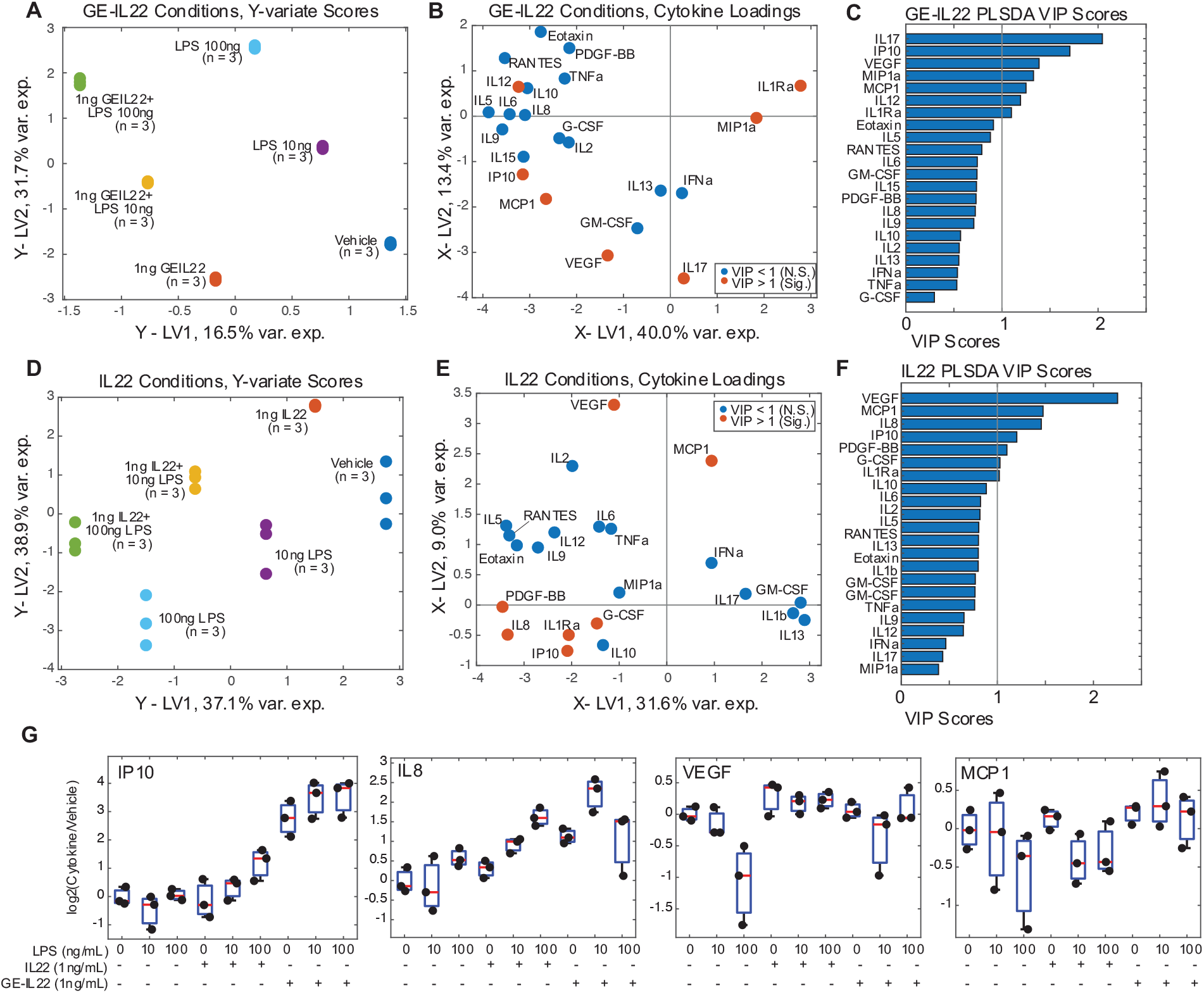
Comparison of Cytokine Responses to Microbial-produced IL22. (A) Y-variate scores for experimental conditions separating GE-IL22 and LPS treatments. (B) Loadings for cytokines in the GE-IL22 PLS-DA model. (C) VIP scores for cytokines in GE-IL22 PLS-DA model. (D) Y-variate scores for experimental conditions separating IL22 and LPS treatments. (E) Loadings for cytokines in the IL22 PLS-DA model. (F) VIP scores for cytokines in IL22 PLS-DA model. (G) Barplots of representative cytokines from core IL22-responsive signature.

The germ-free gut model was challenged by 10 uM and 100 uM of LPS alone and in combination with 1ng/mL of mouse IL-22 and 1ng/mL of our microbe-produced IL-22, here termed “GE-IL-22” for (GE: genetically engineered). We profiled cytokine secretion by the gut model using a 27-plex cytokine assay on a BioPlex-3D Luminex system. To identify those cytokine signals most strongly associated with the inflammatory stimulus (LPS) and anti-inflammatory therapeutic (IL-22 or GE-IL-22) we constructed two Partial Least Squares Discriminant Analysis (PLS-DA) models, one for the GE-IL-22 conditions and one for the pure IL-22 stimulation controls, to relate cytokine concentrations (X-block) to a 2-dimensional Y-block consisting of LPS concentration [0ng, 10ng, 100ng] and IL-22 treatment status [0, 1]. The cytokines most predictive of inflammatory stimulus and IL-22 treatment were identified via variable importance of projection (VIP) score analysis and analysis of cytokine loading coefficients.

For the GE-IL-22 treatments, we identified two axes of separation in the Y-variate block of the PLS-DA model, with GE-IL-22 treatment associated with negative scores on LV1 and increasing concentration of LPS treatment associated with positive scores on LV2 (Figure 6A). Seven cytokines significantly predicted separation between experimental conditions, with IL1Ra, MIP1a, and IL12 associated with escalating LPS concentration (Figure 6B,C). Changes in VEGF, IP10, IL17, and MCP1 were more strongly associated with GE-IL-22 treatment, indicating that these form the core response cytokines of our circuit.

For the pure IL-22 control treatments, we identified the two axes of separation in the Y-variate block of the PLS-DA model separating inflammatory stimulus, more negative scores on both LV1 and LV2 associated with higher LPS concentrations, and more positive LV2 scores associated with IL-22 treatment (Figure 6D). As with the GE-IL-22 treatment, seven cytokines were significantly predictive of response to pure IL-22, with four cytokines (VEGF, IP-10, MCP-1, IL-1Ra) in common with the GE-IL-22 condition and 3 (PDGF-BB, G-CSF, IL-8) being unique to the pure IL-22 conditions (Figure 6E, F). VEGF was the most discriminatory cytokine in the pure IL-22 conditions and ranked in the top 3 for the GE-IL-22 conditions, suggesting VEGF signaling is the core response for IL-22 stimulation in our intestinal model.

As a complementary analysis to the global cytokine analysis via PLS-DA, we also constructed per-cytokine generalized linear models (GLM) predicting cytokine concentration as a function of LPS concentration, IL-22 stimulation, and an interaction term between the co-stimulation terms for both the GE-IL-22 and pure IL-22 stimulation experiments (Tables S3 and S4). In the GE-IL-22 models, seven cytokines were significantly associated with the experimental conditions, with four (VEGF, IL-17, IP-10, IL-1Ra) overlapping with the multivariate PLS-DA model and three (IL-8, IL-6, IL-15) being significant only in the GLMs. For the pure IL-22 conditions, only four cytokines (IL-8, VEGF, IP-10, IL-1Ra) were associated with the experimental conditions. Critically, between the GLM and PLS-DA models, we identified a standard bio-active signature of cytokines associated with IL-22 stimulation, both by GE-IL-22 and pure IL-22 (Figure 6G), indicating the microbial therapeutic produces similar biological responses to the pure cytokine, with minimal off-target effects by other bacterial factors.

### 2.7 Discussion

This study presents the engineering and computational modeling of a prototype living theranostic system that combines autonomous local sensing with external manual control. We have successfully designed and implemented an inflammation-responsive AND logic gate within *E. coli* Nissle. This achievement is the first operational AND logic gate that responds to an inflammatory-associated biomarker in probiotic *E. coli* Nissle. Integrating a two-input AND logical gate is crucial in developing immune-responsive therapeutic. In this system, one input corresponds to the clinically relevant inflammation indicator, tetrathionate, while the second input is manually tunable aTc. To optimize the performance of our engineered circuit, we have selected *E. coli* Nissle due to its proven safety profile in human and murine gut applications, rendering it an ideal candidate for microbiome therapeutic endeavors [55].

Additionally, we have comprehensively examined the circuit’s robustness under diverse parameter settings. This investigation aids in refining the selection of genetic components employed within the circuit design. By conducting experimental variations in ribosome binding sites, we have successfully demonstrated that the enhanced response of the tetrathionate two-component system can be replicated by manipulating ribosome binding rates, as affirmed by our computational model.

Computationally, we designed chemical reaction networks that model our microbial-based circuits *in silico*. We screened a wide variety of parameters, drawing from previously published datasets. Our sensing system modeling found that varying ribosome binding rates to regulator mRNA transcripts vary the response and overall sensitivity to the tetrathionate input. Guided by our model, we experimentally optimized the sensitivity of the two-component tetrathionate inflammation system in *E. coli* Nissle. We screened a library containing various RBS strengths and quantified the circuit’s sensitivity as a function of fluorescent readout. Our tetrathionate-responsive circuit demonstrated high dynamic range across therapeutically relevant concentrations of tetrathionate. We found experimentally varying RBS strength results in different fold changes consistent with our model.

We incorporated AND logic sensing in our engineering bacterial chassis by placing inputs of the split activator AND gate under the regulation of the tetrathionate and IPTG response promoters. After experimental tuning to minimize leak, the dynamics of our circuits were activated when both tetrathionate and IPTG inducers were present at maximum induction concentrations. By creating a chemical reaction network model of our complete AND gate, we were able to identify potential causes for leak in our system. Experimentally, these cases may offer interesting pathways for studying protein-based logic gates. As we engineer additional functionality into our circuit, we hope that our *in silico* RBS tuning and insight into leak will offer us more understanding of our *in vitro* results.

We demonstrated successful secretion of IL-22 through the hemolysin pathway for the first time in *E. coli* Nissle, which has downstream applications in anti-inflammatory treatment. We combined our optimized AND gate with IL-22 secretion to show tetrathionate and aTc-dependent secretion of our target protein. Through linking these two aspects of our project, we created a logic gate that can sense inflammation and a secondary input and respond by secreting approximately 700 pg/ml of anti-inflammatory cytokine mouse IL-22.

In summary, our results demonstrate the usefulness of modular synthetic biological parts and circuit components to design circuits in microbial chassis capable of logically combining two independent input signals, one of which is associated with medical applications. We find that logic gates previously described by Wang et al. can be optimized in *E. coli* Nissle, allowing for future directions in OR, NOT, NOR, and NAND integration. We showed that the hemolysin secretion system characterized by Fernandez et al. can secrete IL-22 in *E. coli* Nissle, extending the medical applications in microbiome engineering and using the synthetic biology toolkit.

In future work, we will characterize the biological activity of our AND gate IL-22 secretion output. We will explore our circuit’s ability to sense and respond to medically-induced inflammation and input signal aTc *in vivo*. The engineered circuit’s functional stability moving from a controlled, *in vitro* environment to the gut microbiome’s competitive environment presents a significant challenge from the competition and metabolic burden perspective. As a continuation, we aim to engineer the second input to increase spatial targeting within the gut.

Engineered microbes can deliver effective therapeutics with exquisite spatial and temporal resolution in medically relevant inflammatory conditions. Synthetic biology may offer advantages over traditional chronic inflammation therapies by designing targeted drug delivery to tissues affected by disease rather than risk off-target effects. The models, logic optimization, and cytokine secretion reported here are preliminary steps toward this long-term goal.

## 3 Methods

### 3.1 Plasmid Construction

Plasmid maps and screening methods are shown in Figure 7. Sequences are available as GenBank files on GitHub [link]. Circuit diagram plots were created with DNAplotlib [51]. All constructs were assembled using 3G assembly [33]. Constructs were sequence verified by sequencing (Laragen, Inc. Culver City, CA) after amplification with 3G universal primers. The sequence for TtrS was identified from Addgene PKD227. We obtained sequences for TtrR and pTTR sequences from Addgene plasmids pKD233.7-3. gBlocks with these sequences were ordered from Twist Biosciences and resuspended in the IDTE buffer. Logic gate parts were gifts from Martin Buck & Baojun Wang. HrpS was amplified from pBW213 (Addgene 61435), HrpR was amplified from pBW115 (Addgene 61434), pHrpL was amplified from pBW412hrpL-cIgfp (Addgene 61438). The BSAI cut site was removed from HrpS by Gibson assembly. B0030 and RBSH sequences were obtained from [17] and synthesized by Integrated DNA Technologies, Inc (IDTDNA, Coralville, IA). Secretion plasmids pVDL9.3 and pEHLY were gifts from Luis Angel Fernandez. HlyBD genes were amplified from pVDL9.3 and inserted into vectors compatible with AND gate plasmids. The sequence for IL-22 was obtained from [53] and codon optimized for *E. coli*. IL-22 sequences were ordered as gBlocks from IDT and resuspended in IDTE buffer.

**Figure 7.**
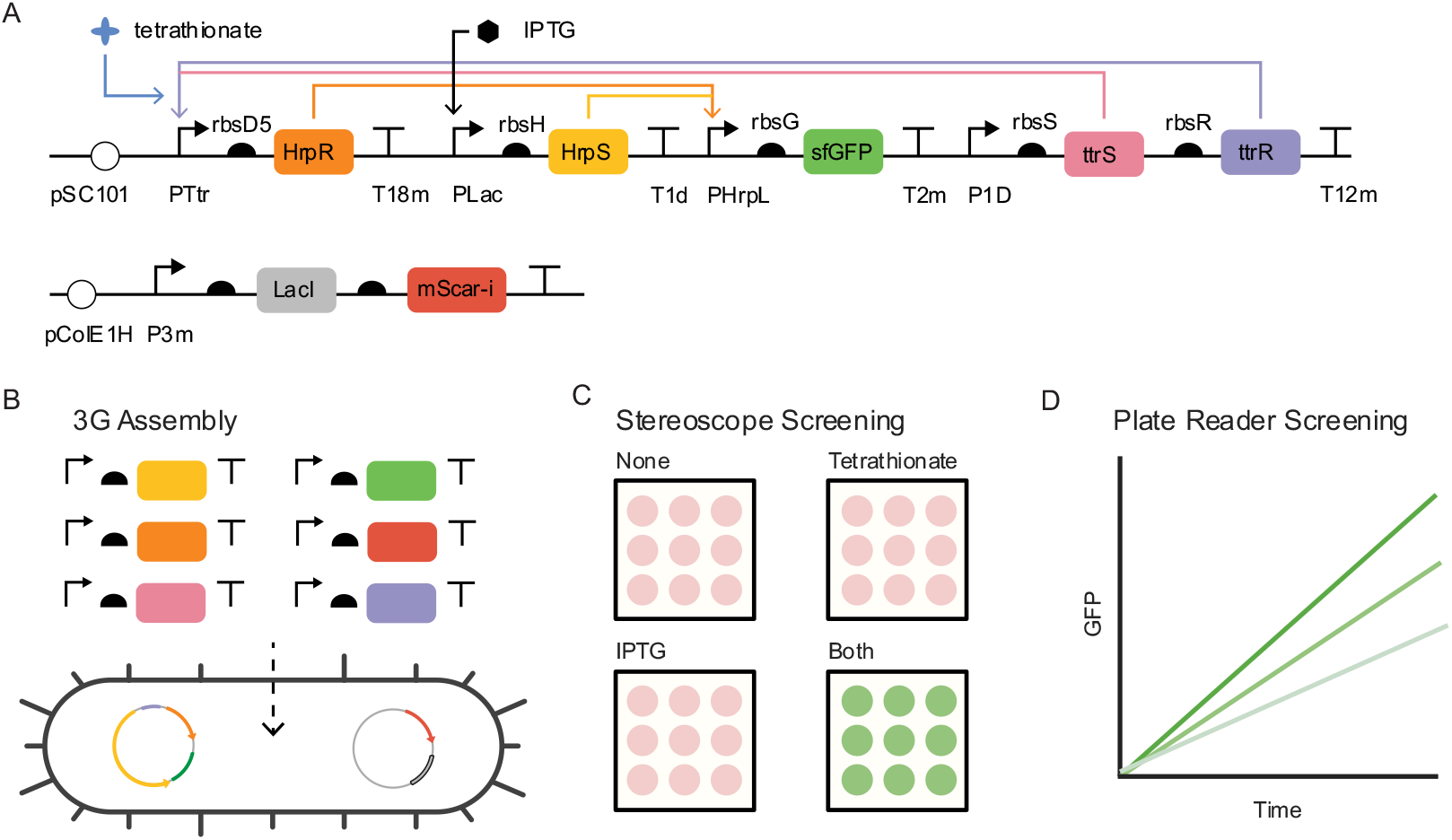
AND Gate Design and Screening. (A) Schematic of engineered logic-based theranostic: We expect GFP expression only when both inducers are present. (B) Golden gate - Gibson Assembly (3G Assembly) method allows for modular assembly of multiple transribtional units for the constribution and transformation of engineered plasmid in probiotic E. coli Nissie 1917. (C) Plate reader assay for Nissie AND constructs. Maximum induction of 1 mM tetrathionate and 1 mM IPTG was used. Fluorescence values were normalized to OD700 readings and readings shown begin at log-growth phase. (D) Plate reader assay for grid of inducer concentrations. Values displayed are GFP fluorescence normalized to OD, divided by the maximal value, achieved at 1 mM IPTG and 1 mM tetrathionate.

### 3.2 Bacterial Strains

Tetrathionate regulator optimization circuits were transformed into chemically competent *E. coli* JM109 (Zymo Research). The AND gate constructs were optimized by making constructs with ribosome binding site library (5’-GAAAGANNNGANNNACTA-3’) in front of regulators in chemically competent Marionette Clo cells prepared from Addgene [40]. Plasmids were miniprepped and re-transformed into electrocompetent Nissle 1917 (Mutaflor). Secretion experiments were performed in *E. coli* HB2151, a gift from Luis Angel Fernandez. Antibiotic concentrations used in all growth were 34 μg/mL chloramphenicol, 100 μg/ml carbenicillin, and 50 μg/mL kanamycin.

### 3.3 *In vitro* Aerobic Experiments

Colonies were screened using stereoscope images of LB agar inducer plates with 1 mM potassium tetrathionate (Sigma Aldrich, catalog no. P2926-25G), 1 mM isopropyl-beta-D-thiogalactoside (Sigma Aldrich, catalog no. I6758), or both. Successful colonies that were only fluorescent in the presence of both inducers were then used for *in vitro* screening. Colonies were grown overnight in M9CA media (Teknova) to saturation. Cultures were then diluted 1:5 and grown for three hours. Characterization was performed in 96 well Matriplates (Dot Scientific, catalog no. MGB0961-1-LG-L). Where applicable, inducer media was prepared in M9CA and diluted outgrown cells. Plates were incubated at 37°C for 23 hours in a Biotek Synergy H2 plate reader with continuous shaking at 282 cpm. Optical densities (OD700) and fluorescence measurements were taken every 5 minutes from the bottom of the plate. GFP excitation and emission wavelengths were 483 nm and 510 nm, respectively. mScarlet-i excitation and emission wavelengths were 565 nm and 595 nm, respectively. Gain 100 was used for both fluorescence channels.

### 3.4 SDS-PAGE

SDS-PAGE (polyacrylamide gel electrophoresis) was used to determine the presence of secreted proteins. 1.5 mL of culture in appropriate inducers was grown overnight at 37°C. Cultures were spun at 8,000 x g for 2 minutes to pellet cells. The supernatant was transferred to a 3 mL syringe and passed through a 0.22 um filter (Pall). According to supplier protocol, the filtered supernatant was then concentrated using 30 kDa centrifugal filters (Amicon). 5 ul LDS sample buffer (Thermo Fisher Scientific) was added to 15 uL of concentrated and incubated at 70°C for 10 minutes. 18 uL was loaded into NuPAGE Bis-Tris Gel (Thermo Fisher Scientific) with MES running buffer (Thermo Fisher Scientific) and run at 200 V for 20 minutes. The SeeBlue Plus2 pre-stained protein standard was used in lane 1 and lane 10 (Thermo Fisher Scientific). Gels were rinsed three times in DI water and then soaked for 1 hr in SimplyBlue SafeStain (Thermo Fisher Scientific) at room temperature with gentle rocking. Gels were destained in deionized water for 2 hours at room temperature with gentle shaking and then imaged on Bio-Rad ChemiDoc MP.

### 3.5 Western Blotting

Western blotting with anti-His antibodies was performed to verify the identity of bands shown in SDS-PAGE gels, except using the Magic Mark XP Western Protein Standard (Thermo Fisher Scientific) as the ladder. Samples were run on an SDS-PAGE gel, as described above. Gels were then rinsed one time in deionized water and placed into an iBlot 2 mini transfer stack (Thermo Fisher Scientific). The transfer stack was loaded onto the iBlot 2 Gel Transfer Device (Thermo Fisher Scientific) and run with the manufacturer’s P0 protocol. After completion of the protocol, the membrane was soaked in Tris-buffered Saline (TBS) pH 7.6 for five minutes at room temperature with gentle shaking. The membrane was soaked for 1 hour in a blocking buffer (3% BSA in TBS). The membrane was soaked for five minutes in TBS and ran overnight on the iBind Western System (Thermo Fisher Scientific) according to the manufacturer’s protocol. To detect His tags on secreted proteins, a Penta His HRP conjugate was used (Qiagen). A Goat anti-Rabbit IgG (H+L) Secondary Antibody conjugated to HRP (Thermo Fisher Scientific) was used to bind to the IgG-tagged ladder. After the binding protocol, the membrane was washed in TBS for five minutes at room temperature with gentle shaking. 20 mL of the SuperSignal West Pico PLUS Chemiluminescent Substrate was made up during the TBS wash. TBS was drained, and the substrate solution was poured over the membrane. The substrate was incubated for five minutes at room temperature with gentle shaking. The membrane was then imaged on Bio-Rad ChemiDoc MP using the Chemi Blot setting. Exposure times ranged from 2 seconds to 3 minutes, depending on the band’s strength.

### 3.6 Germ-Free Gut Model

We cultured Caco-2 (ATCC, passage no. 8) and HT29 (ATCC, passage no.13) colorectal cancer cells separately in Dulbecco’s Modified Eagle Medium (DMEM) containing 10% fetal bovine serum, 1% (v/v) non-essential amino acids, 2 mM l-glutamine, 100 μg/ml streptomycin and 100 U/ml penicillin. At 1x10^5^ cells/cm^2^ concentration, cells were seeded together into the apical chamber of Transwells (Corning Inc, Costar, USA) with a ratio of 9:1 (Caco-2/HT29) for 21 days at 37°C in a carbogen (95% O2, 5% CO2) atmosphere, with the culture medium being changed every two to three days [49,50].

### 3.7 Transwell Stimulation Studies

We challenged the germ-free gut model by introducing 1 uM, 10 uM, and 100 uM of LPS alone and in combination with 1 ng/mL of mouse IL-22 and 1 ng/mL of our microbe-produced IL-22, here termed “genetically engineered IL-22” (GE-IL-22). We profiled cytokine secretion from the gut model using a 27-plex cytokine assay on a BioPlex-3D Luminex system. The other Transwell was treated with 1 ng/ml Escherichia coli O111:B4 lipopolysaccharides on day 22 and harvested on day 23. Cell morphology agreed with the previous description of the Caco-2/HT29 monolayers.

### 3.8 Modeling of Host Cytokine Responses

We constructed PLS-DA models to predict a 2-dimensional Y-block of LPS and IL-22 concentrations [LPS, IL-22] from an X-block of cytokine concentrations, with one model being constructed for the GE-IL-22 stimulations and one for the pure IL-22 stimulations. Models were subjected to 8-fold cross-validation, and up to 8 latent variables were assessed via mean squared error and percent variance explained. For the GE-IL-22 and pure IL-22 conditions, a 3-LV model explaining more than 70% variance and producing local minima of MSE were selected to carry forward. Cytokine VIP scores were calculated using the formula from [54], where a VIP score greater than 1 indicates a cytokine is significantly predictive of the sample projections on Y-variate LVs.

For univariate cytokine analysis, we fit a generalized linear model for each cytokine with the concentration modeled as a function of experimental conditions:

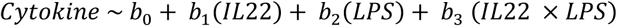

Models were computed using the *plsregress* and *fitglm* functions in MATLAB_2022b, and all code and data necessary to reproduce the results are deposited on the Brubaker Lab GitHub [[link](https://github.com/BrubakerLab)].

### 3.9 Enzyme-Linked Immunosorbent Assay Enzyme-Linked Immunosorbent Assay

We performed an ELISA for quantitatively detecting the genetically engineered production of IL-22. We induced an overnight culture of the engineered strain LM83 with 1 uM concentration of aTc (Sigma Aldrich, catalog no. 37919) and 1 uM of tetrathionate (Sigma Aldrich, catalog no. P2926-25G). We collected the culture’s supernatant at 6-hour, 12-hour, and 24-hour time points post-induction and centrifuged the supernatant at 1650 G for 10 minutes, followed by filtration through a 0.2 mm syringe filter. We quantified the concentration of IL-22 using an Invitrogen Mouse IL-22 ELISA Kit (Invitrogen, catalog no. BMS2047).

### 3.10 Model Simulations

We used the BioCRNpyler [38] Python toolbox to generate a chemical reaction network model for the tetrathionate and AND gate systems shown in figures 2D and 3D respectively. Model reactions and parameters are shown in Table S1 and S2. Simulaions were performed using the Bioscrape [39], a sequential Gillespie’s algorithm for each reaction iteration. All code regenerating the simulations is publicly available on Github [[link](https://github.com/lianamerk/Logical_Inflammation/tree/master)].

## 4 Abbreviations

(ARL): Anderson library of ribosome binding sites
(aTc): anhydrotetracycline
(TtrR): cytoplasmic response regulator
(DI): deionized
(DMEM): Dulbecco’s Modified Eagle Medium
(GLM): generalized linear model
(GE-IL): genetically engineered interleukin
(3G): Golden-gate Gibson
(GFP): green fluorescence protein
(G-CSF): granulocyte colony-stimulating factor
(hrp): hypersensitive response and pathogenicity
(IP): Immunoprecipitation
(IBD): Inflammatory bowel disease
(IL): interleukin
(IDT): Integrated DNA Technologies, Inc
(IPTG): isopropylthiogalactoside
(kDa): kilo-Dalton
(LV): latent variable
(LPS): lipopolysaccharide
(TtrS): membrane-bound sensor histidine kinase
(mRNA): messenger RNA
(μg): micrograms
(μL): microliters
(mL): milliliters
(MSE): mean squared error
(ng): nanograms
(PLS-DA): Partial Least Squares Discriminant Analysis
(PCR): polymerase chain reaction
(RBS): ribosome binding site
(SDS-PAGE): sodium dodecyl sulfate–polyacrylamide gel electrophoresis
(TBS): Tris-buffered Saline
(IDTE): Tris-ethylenediaminetetraacetic acid buffer
(T1SS): type I secretion system
(VEGF): vascular endothelial growth factor
(VIP scores): variable importance and prediction

## 5. Author Information

L.N.M.: Conceptualization, construction of tetrathionate and AND gates, investigation, simulation of tetrathionate and AND gates, visualization, analysis, writing – original draft, writing – review and editing.

A.S.S.: Construction of tetrathionate and AND gates, investigation, simulation of tetrathionate and AND gates, analysis.

S.J./J.M./D.K.B.: Investigation, transwell in vitro model, comparison of cytokine response, visualization, analysis, writing – review, and editing.

R.M.M.: Supervision, funding acquisition, writing – review, and editing.

L.N.G.: Conceptualization, supervision, funding acquisition, visualization, analysis, writing – original draft, writing – review, and editing.

## 6 Acknowledgments

We thank William Poole, Ayush Pandy, John Marken, Andy Halleran, Reed McCardell, Mark Prator, Chelsea Hu, Prof. Boo Tsang for technical support, and all other Murray Lab and Tseng Lab members for insightful discussions. We thank Henry Schreiber and Sarkis Mazmanian for their collaboration and discussions regarding this project. The Caltech NSF AGEP Fellowship and Rosen Fellowship support Leopold Green. We thank Caltech CEMI for supporting this work’s future directions. Justin Bois has provided excellent discussions regarding data analysis and availability. Martin Buck and Baojun Wan provided logic gate strains. Julie Haseman (Purdue University) designed figures; others were created with BioRender.com and modified in Adobe Illustrator.

