## Supplementary Information for "Diagnostic and Therapeutic Microbial Circuit with Application to Intestinal Inflammation"

In this section, we describe the two-component system model in detail. The notation $x : y$ denotes a complex between the chemical species $x$ and $y$. Moreover, all binding rates are denoted with a "b" superscript, and unbinding reactions are denoted with a "u" superscript. A superscript $P$ denotes phosphorylated species, and an asterisk superscript denotes an activated form of a species. The model description is given in Table S1, and the parameter values can be found in Table S2. All code regenerating the simulations is publicly available on Github [link].

Table S1: The Two-Component System Model

| Description | Reaction |
|---|---|
| **Transcription and Translation of Regulators** | |
| RNA Polymerase binds to P1d | $\mathrm{P} + \mathrm{P1d} \underset{k_1^u}{\overset{k_1^b}{\rightleftharpoons}} \mathrm{P1d{:}P}$ |
| Transcription of ttrS, ttrR | $\mathrm{P1d{:}P} \xrightarrow{k_{tx}} \mathrm{ttrS_T} + \mathrm{ttrR_T} + \mathrm{P1d} + \mathrm{P}$ |
| Translation $\mathrm{ttrS_T}$ to ttrS | $\mathrm{ttrS_T} + \mathrm{R} \underset{k_2^u}{\overset{k_2^b}{\rightleftharpoons}} \mathrm{ttrS_T{:}R} \xrightarrow{k_{tl}} \mathrm{ttrS_T} + \mathrm{R} + \mathrm{ttrS}$ |
| Translation $\mathrm{ttrR_T}$ to ttrR | $\mathrm{ttrR_T} + \mathrm{R} \underset{k_3^u}{\overset{k_3^b}{\rightleftharpoons}} \mathrm{ttrR_T{:}R} \xrightarrow{k_{tl}} \mathrm{ttrR_T} + \mathrm{R} + \mathrm{ttrR}$ |
| **Tetrathionate Regulator Phosphorylation Pathway** | |
| Tetrathionate (tt) triggering ttrS phosphorylation | $\mathrm{ttrS} + \mathrm{tt} \underset{k_4^u}{\overset{k_4^b}{\rightleftharpoons}} \mathrm{ttrS}^P + \mathrm{tt}$ |
| ttrR binding to unphosphorylated ttrS | $\mathrm{ttrR} + \mathrm{ttrS} \underset{k_5^u}{\overset{k_5^b}{\rightleftharpoons}} \mathrm{ttrR{:}ttrS}$ |
| ttrR binding to phosphorylated ttrS | $\mathrm{ttrR} + \mathrm{ttrS}^P \underset{k_6^u}{\overset{k_6^b}{\rightleftharpoons}} \mathrm{ttrR{:}ttrS}^P$ |
| Phosphorylation of ttrR | $\mathrm{ttrR{:}ttrS}^P \underset{k_7^u}{\overset{k_7^b}{\rightleftharpoons}} \mathrm{ttrR}^P + \mathrm{ttrS}$ |
| Dephosphorylation of ttrR | $\mathrm{ttrR}^P \xrightarrow{k_{dephos}} \mathrm{ttrR}$ |
| **Response Regulator Gene Activation** | |
| Dimerization of $\mathrm{ttrR}^P$ | $\mathrm{ttrR}^P + \mathrm{ttrR}^P \underset{k_8^u}{\overset{k_8^b}{\rightleftharpoons}} \mathrm{ttrR}_2^P$ |
| $\mathrm{ttrR}^P$dimer binding to pTtr promoter | $\mathrm{ttrR}^P + \mathrm{pTtr} \underset{k_9^u}{\overset{k_9^b}{\rightleftharpoons}} \mathrm{pTtr}^*$ |
| RNA Polymerase binds to $\mathrm{pTtr}^*$ | $\mathrm{P} + \mathrm{pTtr}^* \underset{k_{10}^u}{\overset{k_{10}^b}{\rightleftharpoons}} \mathrm{pTtr}^*{:}\mathrm{P}$ |
| GFP Transcription | $\mathrm{pTtr}^*{:}\mathrm{P} \xrightarrow{k_{tx}} \mathrm{pTtr}^* + \mathrm{P} + \mathrm{GFP_T}$ |
| GFP Translation | $\mathrm{GFP_T} + \mathrm{R} \underset{k_{11}^u}{\overset{k_{11}^b}{\rightleftharpoons}} \mathrm{GFP_T{:}R} \xrightarrow{k_{tl}} \mathrm{GFP_T} + \mathrm{R} + \mathrm{GFP}$ |
| **Degradation Reactions** | |
| $\mathrm{ttrS_T}$ and ttrS degradation | $\mathrm{ttrS_T} \xrightarrow{\delta} \varnothing \qquad \mathrm{ttrS} \xrightarrow{\delta} \varnothing$ |
| $\mathrm{ttrR_T}$ and ttrR degradation | $\mathrm{ttrR_T} \xrightarrow{\delta} \varnothing \qquad \mathrm{ttrR} \xrightarrow{\delta} \varnothing$ |
| $\mathrm{GFP_T}$ and GFP degradation | $\mathrm{GFP_T} \xrightarrow{\delta} \varnothing \qquad \mathrm{GFP} \xrightarrow{\delta} \varnothing$ |

Table S2: The Two-Component System Parameters

| S.no. | Param. | Description | Estimate | Ref. |
|---|---|---|---|---|
| 1 | $k_{tx}$ | Transcription rate (transcripts/second) | 0.1 | Estimate |
| 2 | $k_{tl}$ | Translation rate | 1 | Estimate |
| 3 | $k_{dephos}$ | Dephosphorylation rate | 50 | [56] |
| 4 | $k_1^b$ | Binding of RNA polymerase to P1d | 10 | [57] |
| 5 | $k_1^u$ | Unbinding of RNA polymerase and P1d complex | 0.0001 | [57] |
| 6 | $k_2^b$ | Binding of ttrS transcript to its ribosome | 0.3 | [56] |
| 7 | $k_2^u$ | Unbinding of ttrS and ribosome complex | Varies | Estimate |
| 8 | $k_3^b$ | Binding of ttrR transcript to its ribosome | 0.3 | [56] |
| 9 | $k_3^u$ | Unbinding of ttrR and ribosome complex | Varies | Estimate |
| 10 | $k_4^b$ | Binding of tetrathionate to ttrS | 1.6 | [56] |
| 11 | $k_4^u$ | Unbinding of tetrathionate and ttrS complex | 0.016 | [56] |
| 12 | $k_5^b$ | Binding of ttrR to ttrS | 0.0001 | [56] |
| 13 | $k_5^u$ | Unbinding of ttrR and ttrS complex | 6 | Estimate |
| 14 | $k_6^b$ | Binding of ttrR to phosphorylated ttrS | 0.0001 | [56] |
| 15 | $k_6^u$ | Unbinding of ttrR and $\mathrm{ttrS}^{\mathrm{P}}$ complex | 1 | Estimate |
| 16 | $k_7^b$ | Forward rate for phosphorylation of ttrR | 1 | [56] |
| 17 | $k_7^u$ | Reverse rate for phosphorylation of ttrR | 1 | [56] |
| 18 | $k_8^b$ | Dimerization rate of phosphorylated ttrR | 0.0083 | [56] |
| 19 | $k_8^u$ | Unbinding of the dimerized complex of phosphorylated ttrR | 0.5 | [56] |
| 20 | $k_9^b$ | Forward rate of activation of pTtr promoter | 0.3 | [56] |
| 21 | $k_9^u$ | Reverse rate of activation of pTtr promoter | 0.0001 | Estimate |
| 22 | $k_{10}^b$ | Binding of RNA polymerase to activated pTtr promoter | 10 | [57] |
| 23 | $k_{10}^u$ | Unbinding of RNA polymerase and activated pTtr promoter complex | 0.0001 | [57] |
| 24 | $k_{11}^b$ | Binding of GFP transcript to its ribosome | 100 | Estimate |
| 25 | $k_{11}^u$ | Unbinding of GFP transcript and ribosome complex | 10 | Estimate |

**Sensitivity Analysis of Two-Component System Model**

We performed a sensitivity analysis of the two-component system model that describes the tetrathionate sensor circuit. We looked at the sensitivity of the output GFP fluorescence to changes in reaction rate parameters. We used COPASI [58], an analysis software for biological models, to compute the normalized sensitivity coefficients of GFP with different parameter values. A sensitivity coefficient is given as the partial derivative of the output for each parameter,

$$S = \frac{\partial[\mathrm{GFP}]}{\partial\theta}, \tag{1}$$

where $\theta$ is any model parameter. The COPASI sensitivity analysis tools compute this using the finite difference numerical approximation. We plot the sensitivity analysis results at the end of the time course to show the effects of each parameter on the steady-state reporter protein level.

From the results shown in Figure S1, we can see that some sensitive parameters are the ribosomebinding and unbinding parameters. Although the model has additional sensitive parameters, they are not all tunable. This simple model-based analysis confirms our hypothesis and results in Figure 2, where we tuned the ribosome binding strength to optimize the two-component sensor performance.

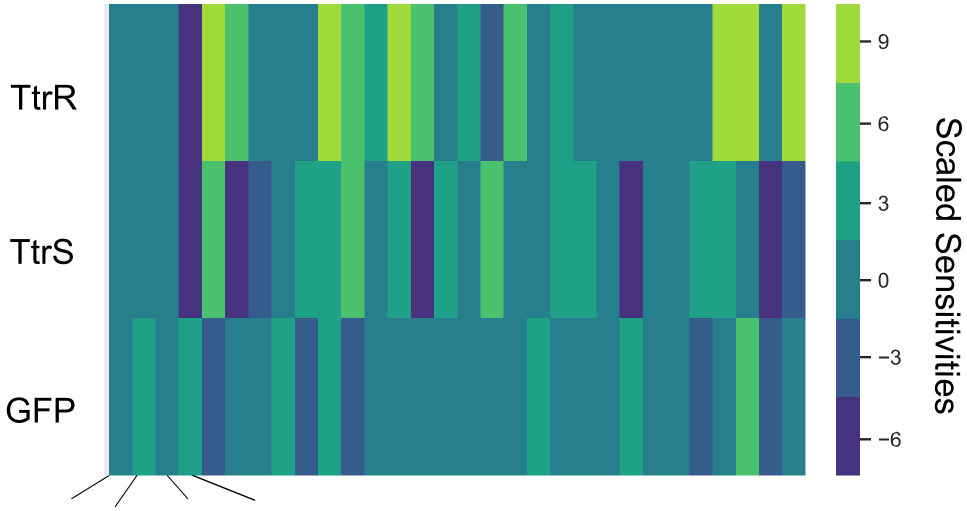


$k^b_2$ $k^u_2$ $k_3{}^b$ $k_3{}^u$ Other model parameters

RBS parameters

**Figure S1.** Sensitivity analysis of tetrathionate sensor model. Heatmap denoting the sensitivity of three key species in the tetrathionate sensor model with respect to model parameters. The tunable parameters in the experiment are the ribosome binding strengths denoted in the Figure.

**Design space exploration with an AND gate model**

In this section we modeled the AND gate circuit by describing the signaling mechanisms and splitactivator genetic components discussed in the previous section. In our model, we explore two key design aspects of the system: (1) the effect of RNA polymerase binding selectively to the combinatorial promoter $P_{HrpL}$ in the presence of HrpR and HrpS proteins and (2) the effect of initial conditions of the LacI repressors. Exploring the design space of the split-activator system guides additional modes for circuit optimization that may prove to be essential when engineering robust circuits for gut-relevant therapeutic applications.

**Effects of RNAP binding**

As shown in Figures S2A and S2B, RNA polymerase can bind to the combinatorial promoter, $P_{HrpL}$, either when both the activators HrpR and HrpS are present, versus when only one of the activators is bound to the promoter. Of course, this binding specificity is not experimentally tunable and depends on the designed constructs and microbial chassis. We model the corresponding reactions in a coarse-grained model to study all of these possible interactions and their effects. We model the logic gate circuit with a non-zero count of the split activator proteins, HrpR and HrpS (R and S, respectively). We performed parameter searches using this logic gate model to study the combinatorial promoter dynamics.

Due to their high sequence similarity, a single hrp activator (R or S) can activate "leaky" transcription, perhaps through mutations in binding domains. To account for this case, our model allows a single activator, DNA:RNA polymerase complex, to activate transcription at a lower rate than the ideal two-activator complex. In Figure 5C, we simulate this effect by varying the unbinding rate of RNAP to the $P_{HrpL}$:HrpR complex by more than 100-fold. We begin with a very low unbinding rate, signifying that RNAP can more readily transcribe the single activator complex, PHrpL:HrpR. Indeed, we see that even small amounts of HrpR are now sufficient to activate the AND gate. The circuit still activates readily at high HrpS, since the $k_u^{HrpR:HrpS} < k_u^{HrpR}$, conferring a preference for the double activator case, increasing the $k_u^{HrpR}$ 100-fold in Figure 5C results in AND gate functionality. This implies that the AND gate's functionality depends on the RNA polymerase's binding specificity to the combinatorial promoter when both activating signals are present. The experimental result for the AND gate implemented in Nissle, as shown in Figure S2, displays similar performance as in the model simulations in the last panel of Figure 5C. Hence, we can hypothesize about this unknown mechanism of RNAP binding and conclude that it is indeed a specific binding that occurs primarily when both activators are present.

To further analyze the system behavior, we expanded the coarse-grained AND gate model to include the expression of the two activators, HrpR and HrpS. We established earlier that the promoter can initiate transcription when both activators are bound but also can leak on at a slower rate if one or neither activator is bound. The detailed model consisting of this construct's transcription and translation reactions also confirms such dynamics. In addition, using the detailed model, we can simulate and understand the dynamics of the activator transcript and protein levels, as well as the dynamics of the reporter transcript and its expression level. We developed this detailed model using BioCRNpyler [38], and for all model simulations, we used parameters and initial conditions consistent with those found in the *E. coli* literature.

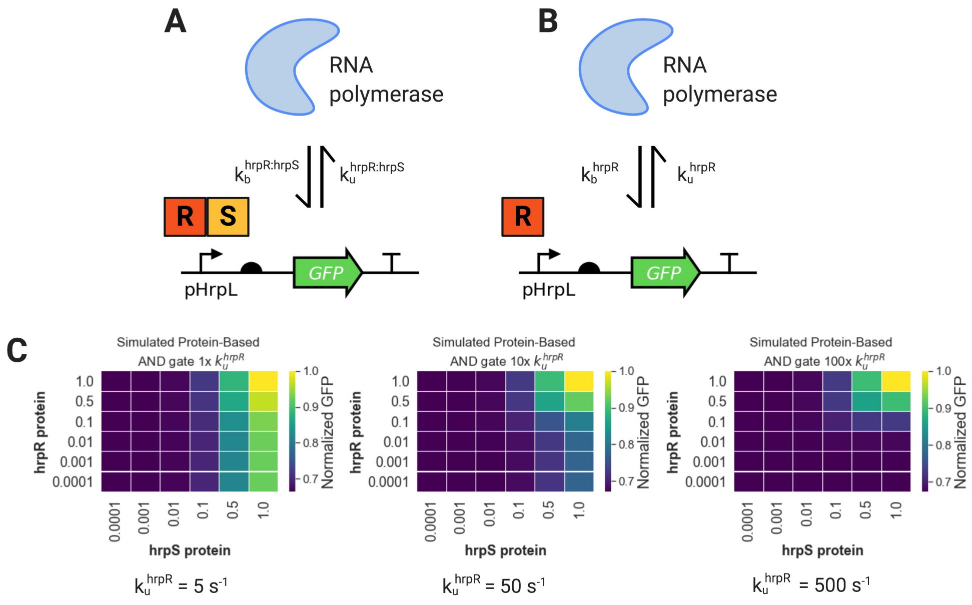


**Figure S2.** Parameter Tuning in Protein AND Gate Model. (A) Both regulators may bind to the promoter region, creating a complex capable of being transcribed. RNA polymerase binds and unbinds with the rates shown. (B) A single activator is also able to trigger transcription. Here, we show the effects of tuning HrpR's ability to trigger transcription. (C) Examples of how decreasing off-rate of RNAP to the $P_{HrpL}$:HrpR affects the output. We see a significant loss of reliance on the second activator, HrpS, when HrpR is allowed to form an active transcription complex that unbinds less frequently. Note, that this result is symmetric if we model HrpS; the AND gate functionality changes in ways consistent with those shown here.

**Effects of repressor initial conditions**

To study the effect of initial conditions for the LacI repressor, we use the LacI repression mechanism we modeled in our full system description; two LacI molecules bind to DNA, forming a complex that represses transcription. A single IPTG molecule can sequester LacI in solution, preventing the repressor complex from binding to DNA in the promoter region. IPTG can also bind to the LacI complex bound to DNA, releasing LacI from the operator region. The AND gate activator HrpR is under the expression of the $P_{Lac}$ promoter. The circuit schematic corresponding to the model is shown in Figure S3A. Note that when LacI is not bound to $P_{Lac}$, HrpR is expressed constitutively. We explore the model simulations by plotting the steady-state GFP values under different input conditions, as shown in the heatmaps in Figure S3B,D.

A key design aspect of this analysis is that if we start with no LacI present, we obtain leaky activation of the AND gate independent of the IPTG levels, as shown in Figure S3B. This is counterintuitive since LacI is constitutively expressed in the system. A possible hypothesis could be that the delay in the transcription of LacI leaves enough time for HrpR expression, even at low levels, hence giving a leaky reporter expression. By looking at the time-course dynamics of the system carefully, we find that this is indeed the case. We observe that there is a sharp spike in actively transcribing $P_{Lac}$. This causes small amounts of HrpR transcription, which is enough to activate the AND gate. It is important to note that increasing the LacI transcription/translation rate and decreasing the PLac transcription rate still need to fix this issue. However, due to bacterial cell division, basal levels of LacI will always be present in the system, diluted from the offspring's parent. Therefore, the case study with zero initial conditions for LacI is not experimentally practical. To account for this in our model, we spiked low concentrations of LacI as an initial condition, thus resulting in a recapitulation of the expected AND gate dynamics, as shown in Figure S3D.

**Engineering AND logic gate in E. coli Nissle**

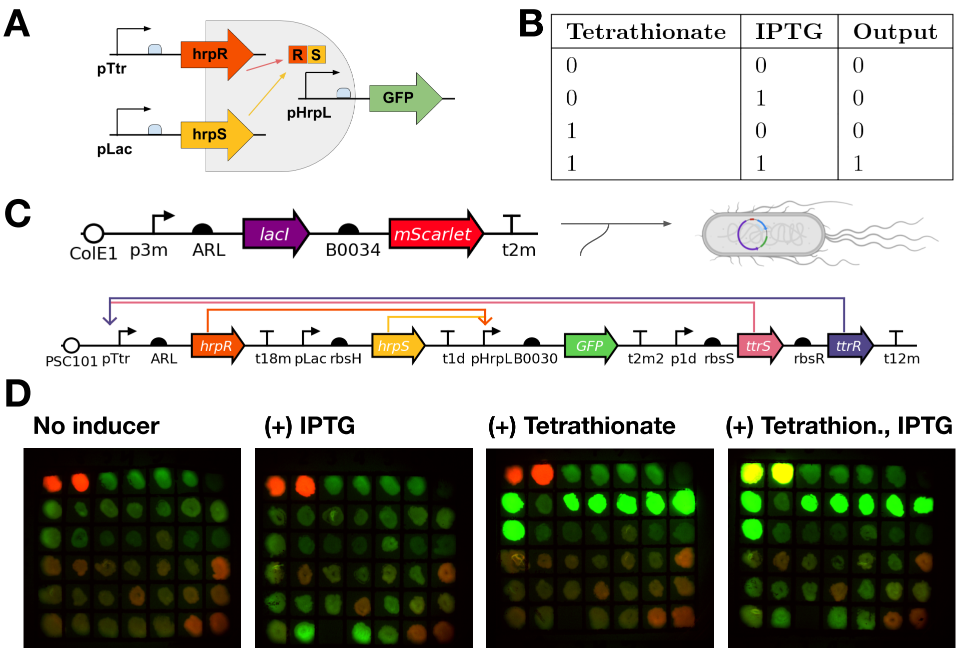


| Tetrathionate | IPTG | Output |
|---|---|---|
| 0 | 0 | 0 |
| 0 | 1 | 0 |
| 1 | 0 | 0 |
| 1 | 1 | 1 |

**Figure S3.** AND Gate Plate Screening: Because many parts of the AND gate required RBS screening for proper strengths without inducing leak, high-throughput plate assays were used to determine which colonies only achieved activation in the dual inducer condition (A) The sigma 54-dependent promoter HrpL is activated by two activators: HrpR and HrpS. The tetrathionate response promoter pTtr drives the expression of HrpR. The second activator, HrpS, is driven by pLac induced by IPTG. Should the AND gate function properly, we expect GFP expression only when both inducers are present. (B) A truth table displaying the expected output. (C.) The circuit diagram for co-transformed plasmids. The low copy pSC101 backbone contains KanR, conferring kanamycin resistance. The high copy pColE1 backbone contains ChlorR, conferring chloramphenicol resistance. (D.) Plate screening of AND gate constructs. LB plates contained max inductions of IPTG (1 mM), Tetrathionate (1 mM), neither, or both. Colonies were streaked on all four plates. Successful colonies were red fluorescent in all plates, signifying constitutive mScarlet production, and green fluorescent on the double inducer plate, signifying AND activation. Colonies in the top left two positions were miniprepped and re-transformed into Nissle.

**Secretion System Optimization**

To increase the yield of any eventual therapeutic output, we explore engineering the well-characterized type I secretion system (T1SS) in our *E. coli* chassis. We incorporated the T1SS by constructing a plasmid containing the HlyB and HlyD sequences and transformed it into electrocompetent tet-inducible HlyA-tagged MalS Nissle cells. We placed the tetR repressor at the end of the inducible construct because Nissle does not contain genome-integrated repressors. We screened a fluorescent version of this construct to identify the proper ribosome binding site strength. We hypothesized that the stoichiometry of HlyB and HlyD is essential for secretion efficiency; thus, we used the ARL RBS pool, proceeding with the HlyB and HlyD genes to screen for optimal expression levels of each secretion component. We used the starch agar plate test to confirm that the secretion pathway was functioning as expected. We observed an increase in cleared (light) areas on the plate, which signifies increased clearance of starch and, thus, an increase in the extracellular secretion of amylase MalS (Figure S4). Interestingly, the control constructs LM42, a Nissle strain without secretion machinery or MalS, does not have high starch clearance. However, the inducible construct does appear to have a larger halo clearance upon MalS activation. While the repressor level of TetR was optimized with a fluorescence screen, there is still a leak in the no aTc condition. Due to the difficulties in interpreting and validating the starch assay qualitative results, we developed a quantitative assay to read out starch concentration.

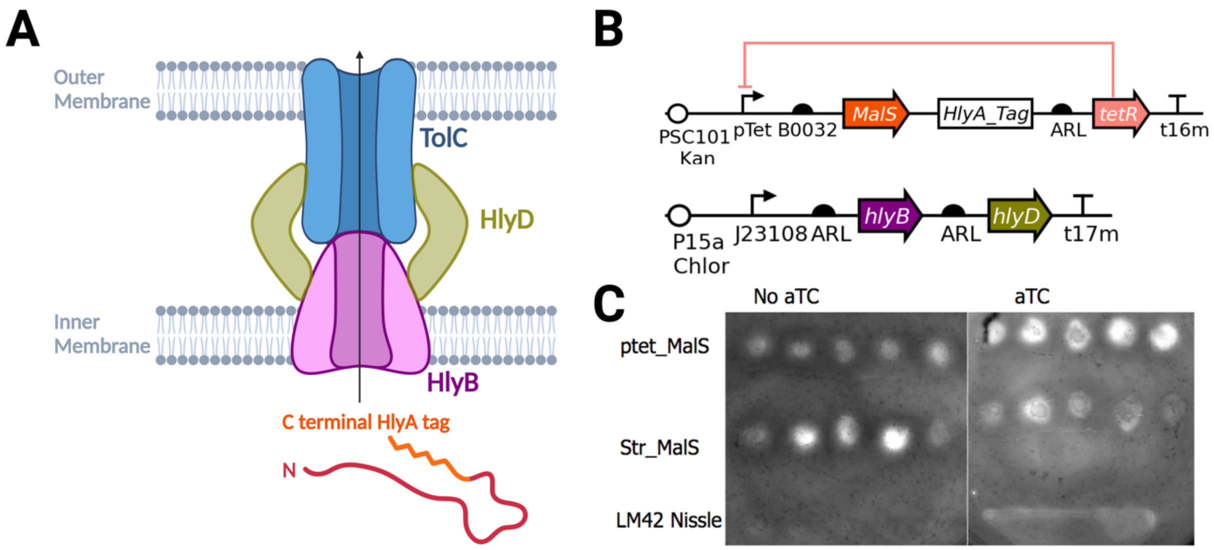


**Figure S4.** HlyBD Secretion System Engineering. (A) The three components of the secretion system are shown with a C-terminal HlyA-tagged protein. (B) The two plasmid secretion systems are used for testing. MalS encodes a starch-degrading enzyme. The 64 amino acid HlyA tag was used. (C) Iodine-stained starch plates with 10 tested colonies. The first row contains aTc inducible MalS, whereas the second is a strong, constitutively expressed MalS. The bottom row is a negative control Nissle cell without secretion machinery or MalS. Adding aTc inducer activates the Tet inducible MalS construct, which has less starch secretion than in the no inducer condition.

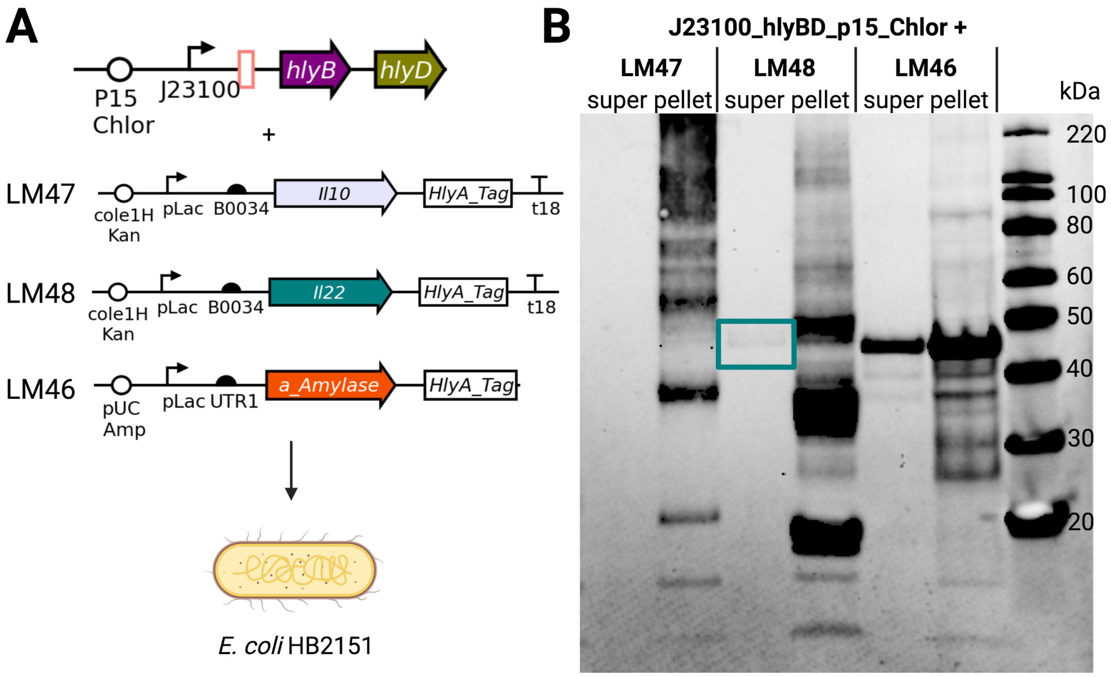


**Figure S5.** Low Yield Anti-Inflammatory Secretion. (A) The successful constitutive secretion machinery construct was isolated and co-transformed with inducible IL-10 or IL-22. (B) A western blot with anti-His antibodies, which bind to N-terminal His tags present on all protein secretion constructs. The expected bands for HlyA-tagged monomers of IL-10, IL-22, and VHH against alpha-amylase are 48 kDa, 46 kDa, and 43.5 kDa, respectively. There are bands of the correct size in the supernatant of both the IL-22 and positive secretion control.

The VHH against alpha-amylase was used as a positive control. While the inducible IL-10 construct failed to show a band of the expected size, inducible IL-22 had a faint band in the supernatant.

We analyzed the bands inside the cell, denoted with "pellet" above the blot shown in Figure S5B. In the IL-10 condition, the three bands are consistent with intracellular IL-10 production. First, the IL-10 monomer without the HlyA tag weighs 19 kDa. The dimer is 38 kDa, and the trimer is 57 kDa. All three of these bands are present internally in the IL-10 samples, leading us to believe the inducible IL-10 construct successfully produces monomeric, dimeric, and trimeric forms of IL-10. The HlyA-tagged IL-10 monomer is expected at 48 kDa. It is unclear from this blot whether a band of that size is present. However, upon further inspection of the interleukin secretion constructs, it was discovered that stop codons were present before the HlyA tag in both IL-10 and IL-22. This is likely the cause for low secretion output in IL-22 and may be the reason there is no external IL-10 seen here. To remove the stop codons from the plasmid, we used Gibson assembly to improve yield in IL-22 and observe secretion in the IL-10 construct.

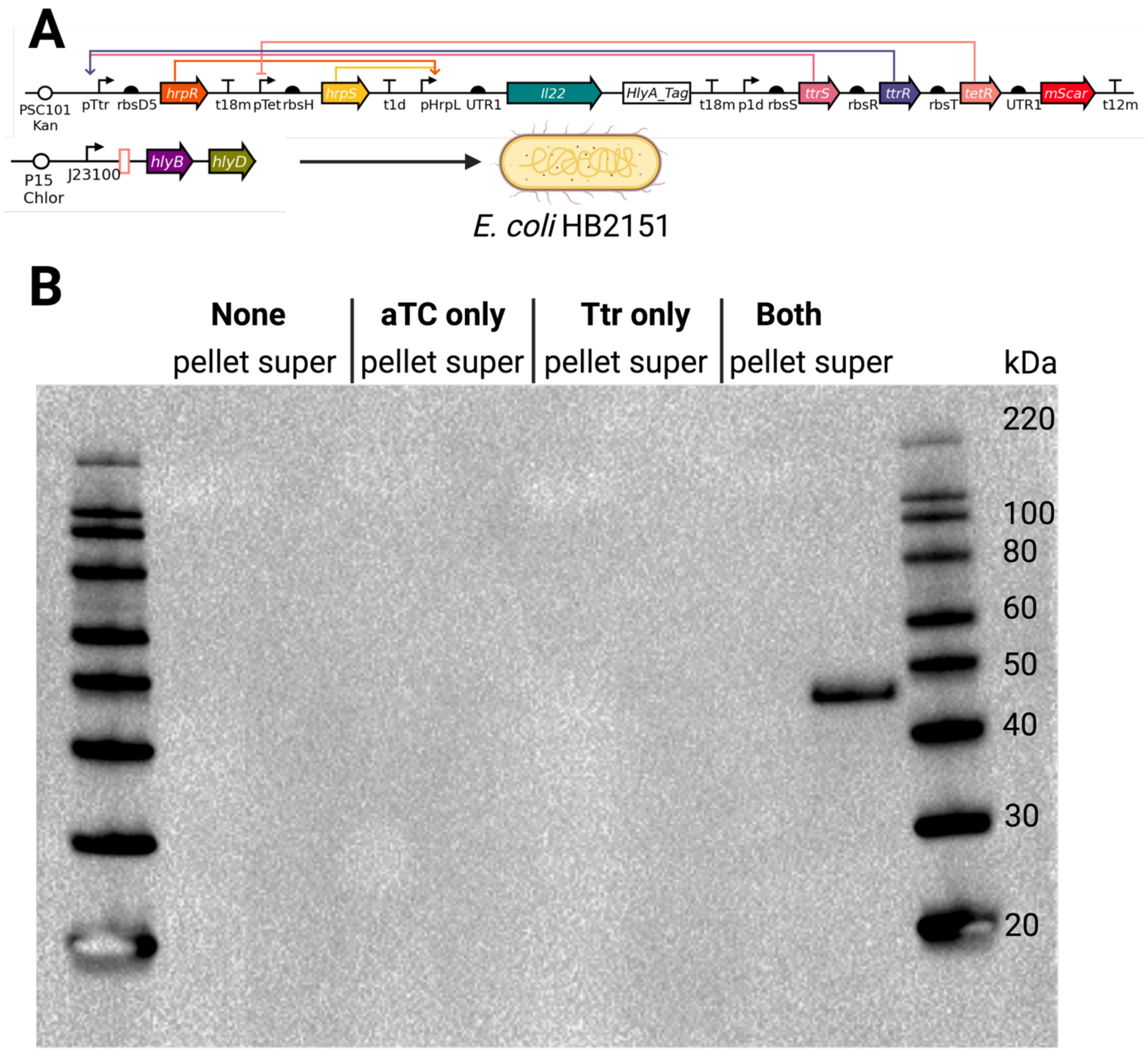


**Figure S6.** AND Gate IL-22 Secretion. (A) AND gate backbone was amplified, and the AND product, driven by pHrpL, was changed from GFP to HlyA-tagged IL-22. Constructs were co-transformed with the constitutive expression hlyBD into *E. coli* HB2151 and sequence verified. (B) A western blot with anti-His antibodies shows IL-22 secretion only when aTC and tetrathionate are present.

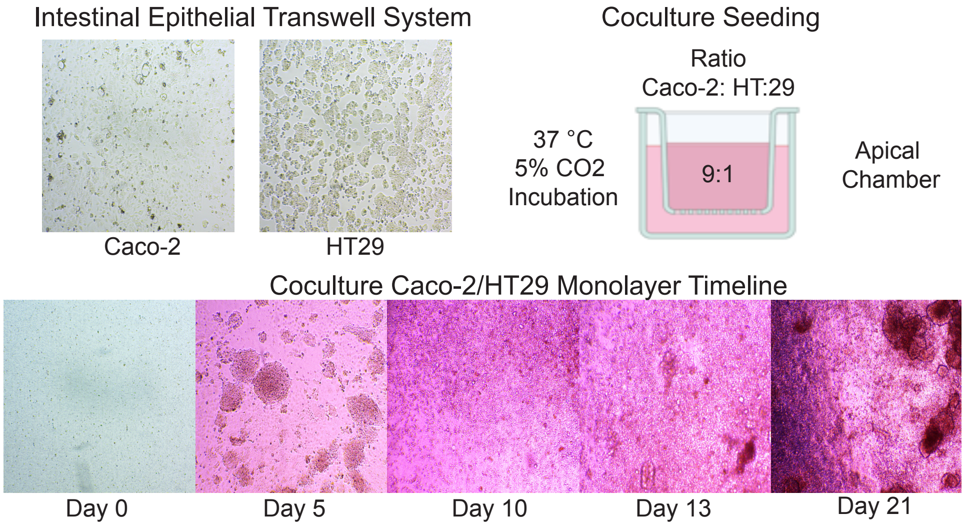


**Figure S7:** Co-cultured Caco2 and HT-29 colorectal cancer cells on transwell inserts to create a germ-free intestinal epithelium mode.
